# Morphological changes of plasma membrane and protein assembly during clathrin-mediated endocytosis

**DOI:** 10.1101/209791

**Authors:** Aiko Yoshida, Nobuaki Sakai, Yoshitsugu Uekusa, Yuka Imaoka, Yoshitsuna Itagaki, Yuki Suzuki, Shige H. Yoshimura

## Abstract

Clathrin-mediated endocytosis (CME) proceeds through a series of morphological changes of the plasma membrane induced by a number of protein components. Although the spatiotemporal assembly of these proteins has been elucidated by fluorescence-based techniques, the protein-induced morphological changes of the plasma membrane have not been fully clarified in living cells. Here we visualize membrane morphology together with protein localizations during CME by utilizing high-speed atomic force microscopy combined with a confocal laser scanning unit. The plasma membrane starts to invaginate ~30 seconds after clathrin starts to assemble, and the aperture diameter increases as clathrin accumulates. Actin rapidly accumulates around the pit and induces a small membrane swelling, which within 30 seconds rapidly covers the pit irreversibly. Inhibition of actin turnover abolishes the swelling and induces a reversible open-close motion of the pit, indicating that actin dynamics are necessary for efficient and irreversible pit closure at the end of the CME.

## Introduction

Cells communicate with the extracellular environment via the plasma membrane and membrane proteins. They transduce extracellular signals and substances into the cellular plasm via cell-surface receptors, channels, and pumps, as well as by various endocytic processes^1–4^. Cells also disseminate their intracellular contents to the extracellular space via exocytosis. These dynamic cellular processes are largely dependent on the assembly and catalytic function of various proteins in the plasma membrane. Clathrin-mediated endocytosis (CME) is conducted by more than 30 different proteins. Extensive studies using fluorescence imaging techniques revealed the spatiotemporal dynamics of individual proteins in living cells^5–7^. In addition, a number of *in vitro* studies revealed unique functions of these proteins in deforming the plasma membrane^8^. For instance, BAR domain proteins bind to the surface of the lipid bilayer and induce membrane curvature and tabulation, and are therefore presumed to be involved in membrane deformation in an early stage of the CME^9^. Dynamin also induces membrane tabulation with a smaller diameter and vesiculation via a nucleotide-dependent conformational change, and therefore has been considered to be involved in vesicle scission process^10,11^.

Despite our increasingly detailed knowledge regarding the cellular dynamics of these proteins *in vivo* and their catalytic activity *in vitro*, the morphological changes of the plasma membrane during CME in living cells have not been studied. This has mainly been due to a lack of imaging techniques for visualizing the membrane. Electron microscopy has made a substantial contribution to the study of CME, owing to its high spatial resolution. The detailed morphological changes of the plasma membrane, together with the assembly of proteins, such as clathrin, have been imaged and analyzed in a series of images to understand the entire process of CME^12–16^. However, aligning a thousand EM snapshots still suffers from a large limitation in the time resolution. In contrast to EM, fluorescence labeling and imaging techniques are powerful tools for studying protein dynamics in living cells. Recent advances in these techniques allow time-lapse imaging of a single protein molecule in a living cell with sub-second time resolution. However, it is not suitable for imaging morphological changes of the plasma membrane in a living cell at a sub-micrometer scale.

Scanning probe microscopies, including atomic force microscopy (AFM), are powerful approaches for characterizing the surface of a specimen at nanometer resolution. Notably, high-speed AFM (HS-AFM) has been utilized to visualize various molecular structures and reactions at sub-second resolution *in vitro*^17–20^. We recently developed a HS-AFM for live-cell imaging and successfully visualized structural dynamics of the plasma membrane in living cells^21,22^. In this study, we utilize this HS-AFM to analyze the morphological changes of the plasma membrane during CME. To understand the role of specific proteins during the morphological change, HS-AFM is combined with confocal laser scanning microscopy (CLSM), so that we could simultaneously visualize membrane structures and protein localizations during CME in living cells. Overlaying AFM and fluorescence images reveals the dynamics of protein assembly and concomitant morphological changes of the plasma membrane with high spatial resolution. Especially, we elucidate the role of actin in the closing step of the CME.

## Results

### Hybrid imaging with combined HS-AFM and CLSM

To reveal protein-induced membrane deformation during CME in a living cell, we first established a hybrid imaging system with HS-AFM and a confocal laser scanning microscope. We previously reported the development of a tip-scanning AFM unit and its combination with an inverted optical microscope with a fluorescence illumination unit^21,22^. In this study, we combined the HS-AFM unit with an inverted optical microscope equipped with a confocal laser scanning unit to increase the optical resolution. To obtain stable imaging, the stage was re-designed. The detailed configuration of the stage is described in Figures 1a and 1b. A cross-shaped movable XY-stage is mounted on the base plate of the inverted optical microscope stage, which allows the specimen to move independently of the AFM unit and the objective lens. The AFM scanning unit now has a 6.0 × 4.5 µm^2^ scanning area, which is larger than the previous unit (4.0 × 3.0 µm^2^), to enable imaging of a larger area of the cell surface.

**Figure 1.**
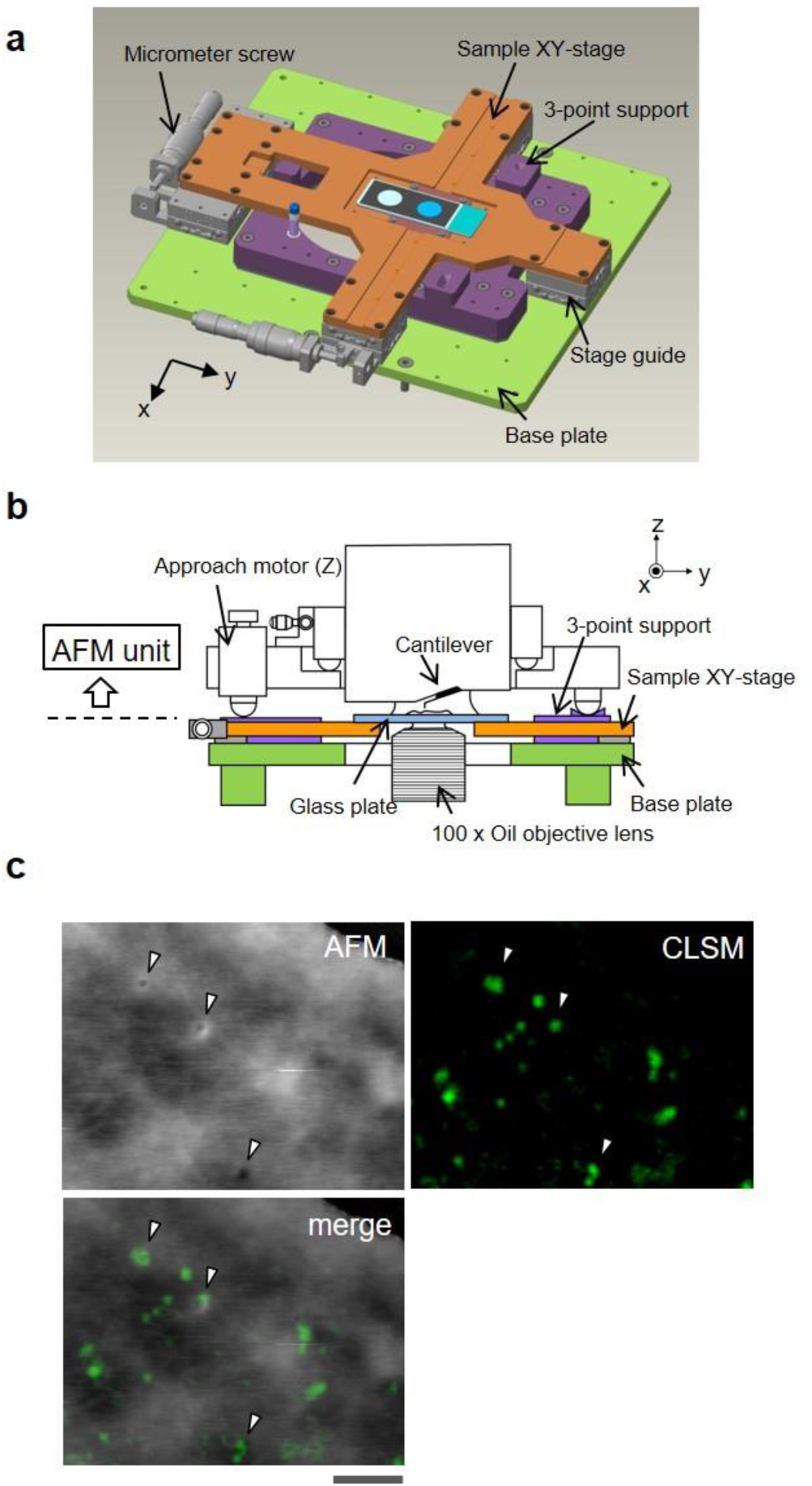
Aligning the confocal image and the AFM image. (**a**) Schematic illustration of the sample stage. A cross-shaped movable XY-stage (orange) is mounted on the base plate (light green) of the inverted optical microscope (IX83) via a stage guide (gray) equipped at each of the four ends of the cross. A 3-point support plate (purple) for mounting the AFM scanner unit is fixed on the base plate with a configuration that does not hinder the sliding of the XY-stage along the x-and y-axis. These setups allow the sample stage to move independently of the AFM unit and the optical axis. (**b**) Side-view of the high-speed AFM unit mounted on the stage illustrated in (**a**). (**c**) Overlaying a confocal image and an AFM image. COS-7 cells expressing EGFP-clathrin were fixed with 5% paraformaldehyde and subjected to AFM (top left) and CLSM (top right) imaging. The x-y position of the probe tip was determined as described in Supplementary Fig. S1. Two images were overlaid (bottom left) based on the x-y center position. Scale bar: 1 µm.

We then established a procedure for aligning confocal fluorescence and AFM images. The details of the alignment method are described in Supplementary Fig S1. In brief, the probe was brought to approach and attach on the glass surface without scanning. The z-position of the probe tip and confocal plane was also aligned by setting the glass surface as a reference position (z = 0). The x-y position of the probe tip on the optical axis was determined by imaging an auto-fluorescence signal of the probe (Supplementary Fig. S1a and S1b). The center position of the probe fluorescence was defined as an origin (x, y = 0, 0), and the x-y position of the AFM image also refers to this scale (Supplementary Fig. S1b). Once the reference position of the probe tip and optical axis was determined, the cross-shaped sample stage allowed the specimen (cell) to move in x-y directions without changing the relative position between the HS-AFM and CLSM.

The spatial accuracy of the hybrid imaging described above was tested by observing a chemically fixed cell. COS-7 cells expressing EGFP-fused clathrin were fixed and observed by hybrid imaging. Several different membrane structures could be identified in the AFM image of the cell surface (Fig. 1): membrane invaginations of different sizes (diameters); ruffle-like short narrow protrusions; and large swellings. When the AFM image was overlaid on the confocal fluorescence image of EGFP-clathrin based on the x-y position alignment, several clathrin spots readily co-localized with membrane invaginations identified in the AFM image (Fig. 1c). The section profile analysis revealed that the diameter of the membrane invaginations (pit) identified in the AFM images ranged from 100 – 400 nm, whereas those co-localized with a clathrin spot ranged between 150 and 400 nm. The aperture size of the clathrin-coated pit (CCP) previously observed by electron microscopy was slightly smaller than our observation (20 – 175 nm)^23^. This is apparently due to the procedure of the image analysis. In the EM image, the narrowest aperture (inner diameter) was measured, whereas the outer diameter was measured in the AFM (The details of the AFM image analysis are provided in Supplementary Fig. S2).

To evaluate the accuracy of the image alignment, the position (x, y) of the membrane invagination in the AFM image was compared with that of the corresponding fluorescence spot in the confocal image. The average distance between two spots was 35±10 nm (deduced from 4 spots in the same image). Considering the diameter of the membrane invaginations (150-400 nm), we concluded that our image overlaying procedure is accurate enough to merge the clathrin spot with the AFM image. It should be noted that the number of membrane invaginations in the AFM image was less than that of the fluorescent clathrin spots: i.e. there were some clathrin spots which did not co-localize with membrane invaginations in the AFM image. Such clathrin signals could be due to either clathrin-coated vesicles (CCVs) that had already budded from the plasma membrane, or CCPs that formed in the basal surface of the cell.

### Live-cell hybrid imaging of membrane morphology and protein assembly in the CME

To follow the entire process of CME, we then performed time-lapse hybrid imaging of living cells. A live COS-7 cell expressing EGFP-fused clathrin was observed by HS-AFM and confocal microscopy in culture medium containing fetal bovine serum (FBS). Both AFM and CLSM images were obtained every ten seconds, and overlaid. Small membrane invaginations as described in Fig. 1 were co-localized with EGFP-clathrin spots over several minutes (Fig. 2a, 2b) (see video in Supplementary Movie S1). The offset of the fluorescence spot from the corresponding membrane pit in the AFM image was less than 60 nm throughout the observation (Supplementary Fig. S3), and no severe drift between two merged images was detected. This indicates that the AFM and optical imaging units are stably combined, so that the relative position of two units is well-maintained during a long period of time-lapse imaging.

**Figure 2.**
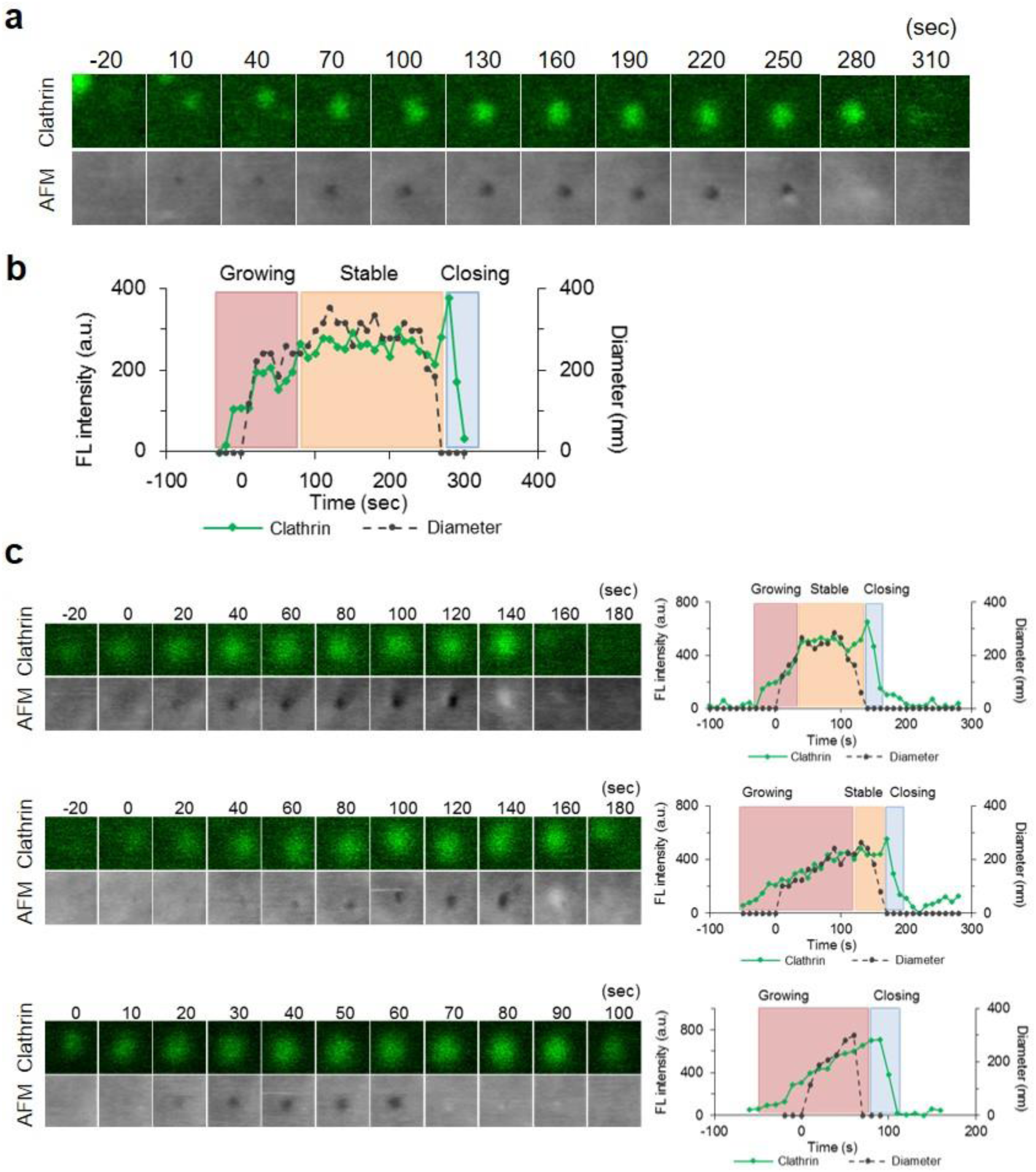
Hybrid AFM imaging reveals morphological changes of clathrin-coated pits in living cells. In culture medium, live COS-7 cells expressing EGFP-Clathrin were subjected to time-lapse hybrid AFM imaging. The position of the probe tip was aligned as described in Supplementary Fig. S1. (**a**) The fluorescence image (top) and AFM image (bottom) of a representative CCP. Image size: 0.6 × 0.6 µm^2^. (**b**) The fluorescence intensity (green) and diameter of the CCPs in the AFM images (dotted black) in (a) are plotted against time. Based on the changes of the clathrin signal, the whole process is divided into a growing phase, a stable phase, and a closing phase. (**c**) Variations of CCPs observed by hybrid time-lapse imaging. Individual CCPs have large variations in the duration of the growing and stable phases.

Morphological changes of the plasma membrane during CME were further analyzed in a series of AFM images together with fluorescence signals from EGFP-fused clathrin. The fluorescence intensity of the clathrin spot and the diameter of the pit in the AFM image were plotted against time (Fig. 2b). The clathrin signal appeared 20-30 seconds before the membrane started to deform. The clathrin signal then increased (growing phase) until it reached a stable phase as demonstrated by a previous study^24^. During the growing phase, the aperture of the pit in the AFM images also increased (Fig. 2b), suggesting that the size of the pit also enlarged during this period of time. During the stable phase, the aperture also remained almost constant. Interestingly, there were large variations in the duration of the growing and stable phases; the growing phase ranged from 40-280 sec, and the subsequent stable phase lasted between 0 and 260 sec (N=17) (Fig. 2c). Following the stable phase, the closing step proceeded over a short period of time (20-50 sec) (Fig. 2b, 2c). Notably, the clathrin spot remained for another 20-30 seconds after the pit closed, and then suddenly disappeared. This could indicate that either the clathrin coat remains on the vesicle and is eventually disassembled by GAK^25,26^, or the vesicle eventually moves out of the focal plane of the CLSM. The total lifetime of the CCP ranged from 40 to 330 sec (N=113).

### Assembly of other CCP-related proteins

Assemblies of other CCP-related proteins were also investigated by time-lapse hybrid imaging. Epsin is known to add bending stress to the lipid bilayer at an early stage of the CME, thus changing the membrane curvature^27^, and it recruits clathrin to the pit surface. COS-7 cells simultaneously expressing mCherry-fused Epsin and EGFP-fused clathrin were subjected to time-lapse hybrid imaging. Epsin started to assemble on the plasma membrane prior to the membrane invagination, which was similar to clathrin (Fig. 3a and 3b; Supplementary Movie S2). However, statistical analysis of the timing of assembly and membrane invagination revealed that epsin assembles prior to clathrin; fluorescence spots of epsin and clathrin appeared 47 ± 9 and 34 ± 11 sec, respectively, before the membrane started to invaginate (Fig. 3e; Supplementary Fig. S4). During the growing phase, fluorescence signals of both proteins increased (Fig. 3b) until the pit closed. Epsin and clathrin signals peaked at ~14 and ~3 sec before the pit closed, and disappeared at ~20 and ~13 sec after the closure, respectively.

**Figure 3.**
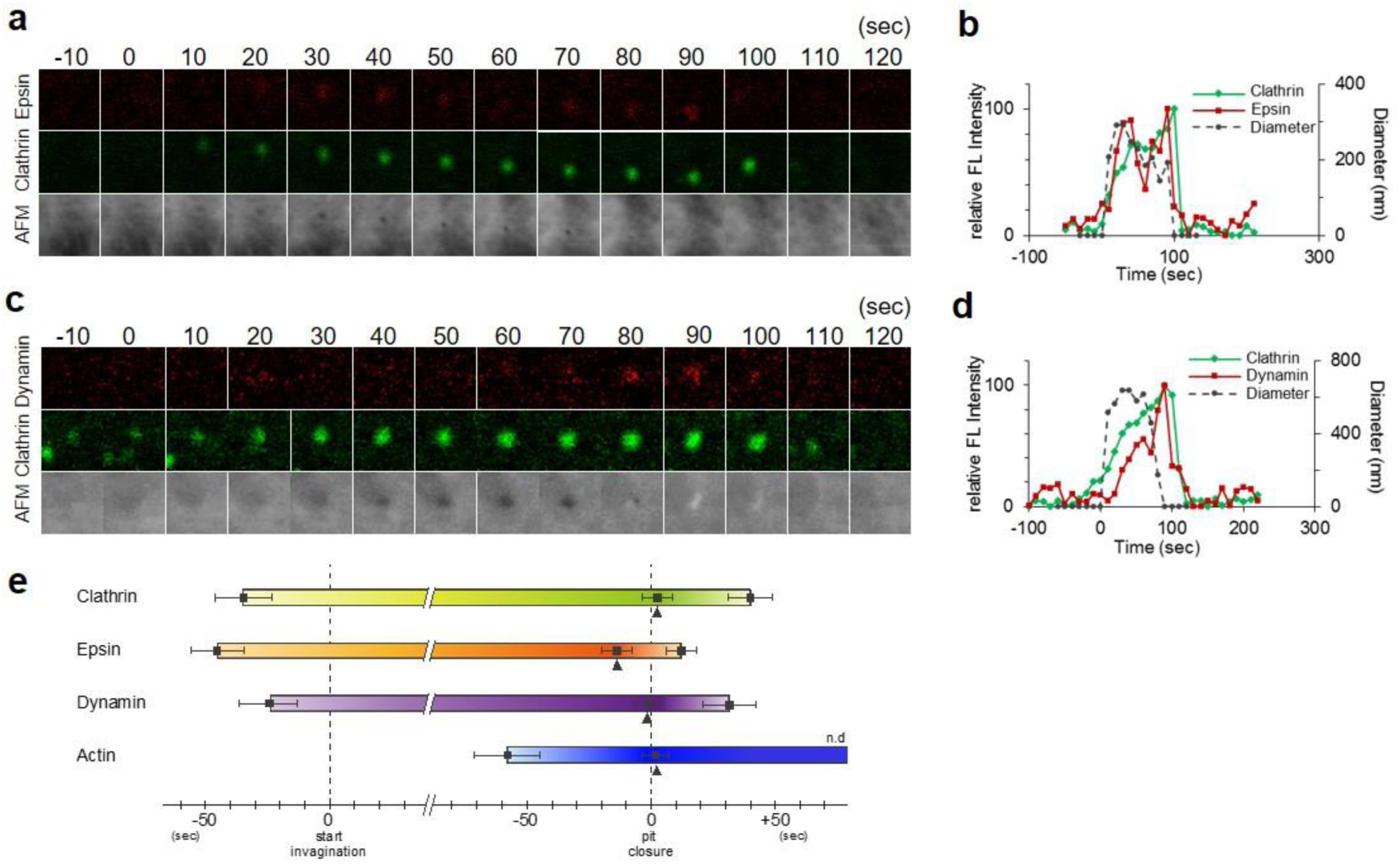
Morphological changes of the plasma membrane and protein assembly during CME. (**a,b**) Time-lapse hybrid imaging of COS-7 cells expressing EGFP-clathrin and mCherry-Epsin. Fluorescence images for epsin and clathrin, as well as AFM images are shown every 10 seconds (**a**). Image size: 1.2 × 1.2 µm^2^. The signal intensities of the fluorescence spots for clathrin (green) and epsin (red), and the diameter of the membrane invagination in the AFM image (dotted black) were plotted against time (**b**). (**c,d**) Time-lapse hybrid imaging of COS-7 cells expressing EGFP-clathrin and mCherry-dynamin. Fluorescence images for dynamin and clathrin, as well as AFM images are shown every 10 seconds (**c**). The signal intensities of the fluorescence spots for clathrin (green) and dynamin (red), and the diameter of the membrane invagination in the AFM image (dotted black) were plotted against time (**d**). (**e**) A summary of protein assembly at CCPs. Based on the AFM images, the time when the plasma membrane started to invaginate (left half) and when the pit had completely closed (right half) were defined as t = 0. The results of hybrid imaging for clathrin, epsin, dynamin, and actin are summarized along this time scale. The timings when the fluorescence signal appeared on the CCP (left half) and disappeared (right half) are plotted. n = 35 for clathrin, n = 8 for epsin, n = 13 for dynamin and n = 14 for actin. Error bars represent S.D.

Dynamin localizes at the neck of the pit and plays a role in the vesicle scission^16^ in the last step of CME. COS-7 cells expressing mCherry-fused dynamin2 (an isoform ubiquitously expressed in a variety of cells) together with EGFP-clathrin were subjected to the time-lapse hybrid imaging. Although dynamin plays a role in the vesicle scission, a dynamin signal started to assemble on the CCP in an early stage of CME as demonstrated in previous studies^5,28^; in our observations, it assembled ~21 sec after clathrin started to appear, and ~25 sec before the membrane invagination began (Fig 3c, 3d, and 3e, Supplementary Movie S3) (see Supplementary Fig. S5 for other observations). The signal gradually increased in the growing phase, but during the stable phase, the dynamin signal was not clearly defined compared to clathrin. The dynamin signal peaked with the same timing as pit closure, and gradually decreased thereafter (it completely disappeared 37.5 sec after the closure), consistent with the notion that it is involved in the last step of CME.

We further confirmed that the membrane pits observed were indeed CCPs and not other types of endocytic structures. Caveolae are found in another endocytic pathway which is mediated by other sets of proteins (caveolin etc.), but also includes invagination of the plasma membrane. mCherry-fused caveolin 1, a major component of caveolae, was expressed in COS-7 cells together with EGFP-fused clathrin, and live cells were subjected to time-lapse hybrid imaging. The clathrin spots did not co-localize with caveolin1 spots during the observation. Overlaying three images (AFM, EGFP-clathrin, and mCherry-caveolin 1) clearly revealed morphological differences between CCP and caveolae (Fig. 4a, see also Supplementary Movie S4). The aperture of caveolae ranged from 80 – 120 nm, whereas that of CCPs ranged 150 – 400 nm (Fig. 4b). Similar to CCP, the aperture of the caveolae observed by AFM was slightly larger than those observed by EM^29^, probably due to the same reason as described above. Caveolae had longer lifetimes than CCPs; the average lifetime of a CCP was 81 ± 55 sec, whereas caveolae remained open for over 400 sec. They also showed different lateral movements in the plasma membrane; the diffusion coefficient of CCPs was 7.3 × 10^−9^ (cm^2^ sec^−1^), whereas that of caveolae was 2.1 × 10^−9^ (cm^2^ sec^−1^) (Fig. 4c, 4d). Taken together, these results indicate that time-lapse hybrid imaging could identify and distinguish the different aperture opening and diffusion kinetics of the two types of invaginations.

**Figure 4.**
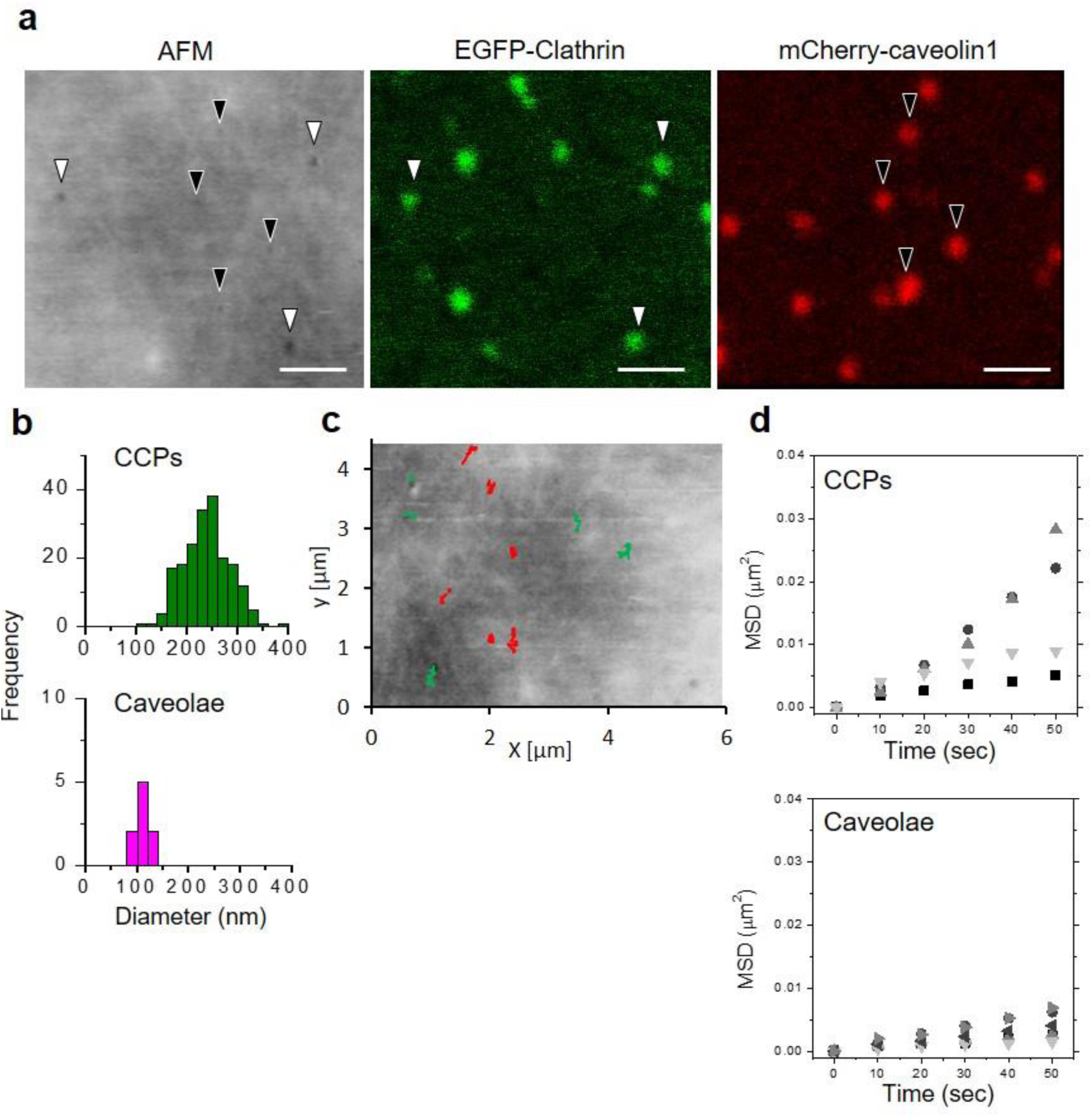
Hybrid AFM imaging distinguishes clathrin-coated pits from caveolae in live cells. COS-7 cells expressing both EGFP-Clathrin and mCherry-caveolin 1 in culture medium were subjected to hybrid time-lapse AFM imaging. (**a**) The x-y alignment of the obtained AFM image (left), EGFP image (middle), and mCherry image (right) was performed as described in Supplementary Fig. S1. Small membrane invaginations which co-localized with an EGFP signal or with an mCherry signal are indicated by black and white arrowheads, respectively. Scale bars: 1 µm. (**b**) The diameters of the membrane invaginations that co-localized with a clathrin signal (top panel) or with a caveolin signal (bottom panel) were measured by section analysis and are summarized as a histogram. (**c,d**) Lateral movements of CCPs and caveolae were analyzed. The trajectories of the pit center (green for clathrin, red for caveolae) were superimposed on the AFM image (**c**), and the mean square displacement (MSD) was plotted against time (**d**). Top panel: CCP, bottom panel: caveolae.

### Unique morphologies of the plasma membrane during CME

In contrast to the growing and stable phases, which continue for more than a minute, the closing and disassembly phase was completed relatively rapidly (< 30 sec). In many cases, the membrane aperture suddenly disappeared (Fig. 2). However, the detailed image analyses revealed several unique membrane structures and dynamics in the closing step of the CCP, which include: i) capping, ii) two-step, and iii) re-opening (Fig. 5a-c). The capping motion was frequently observed in more than 50% of the CME events (Fig. 5d). A small membrane region adjacent to the CCP swelled and eventually covered over the pit (Fig. 5a). The section profile analysis (Supplementary Fig. S2) revealed that the diameter and height of the swelling region was 378 ± 62 nm in diameter and 38 ± 10 nm in height (N=13), respectively, which is comparable to the pit size. The entire closing motion took 23 ± 13 sec (N=48). Two-step closing was observed in 10-20% of the CME events (Fig. 5d). The pit aperture first decreased to ~120 nm, and then disappeared (Fig. 5b). The duration of the small-aperture step was <40 sec. Our trial of AFM imaging with higher time resolution (2 sec frame^−1^) revealed two-step motions with faster small-aperture steps (~10 sec). These results indicate that many CCPs close with a two-step motion, but the duration of the smaller-aperture step varied from several secs to 40 sec. A re-opening motion was also observed in 10-20% of the total CMEs (Fig. 5d). A pit once closed completely then re-opened after several frames of the closure (Fig. 5c). The duration of the closed state varied between 40-70 sec. Interestingly, some CCPs underwent a second close and re-open cycle, suggesting that the pit closure and re-opening is largely reversible. During the re-opening cycle, the clathrin signal decreased when the pit closed and re-increased when the pit re-opened (Fig. 5c), indicating that the aperture of the pit and the amount of clathrin are tightly coupled.

**Figure 5.**
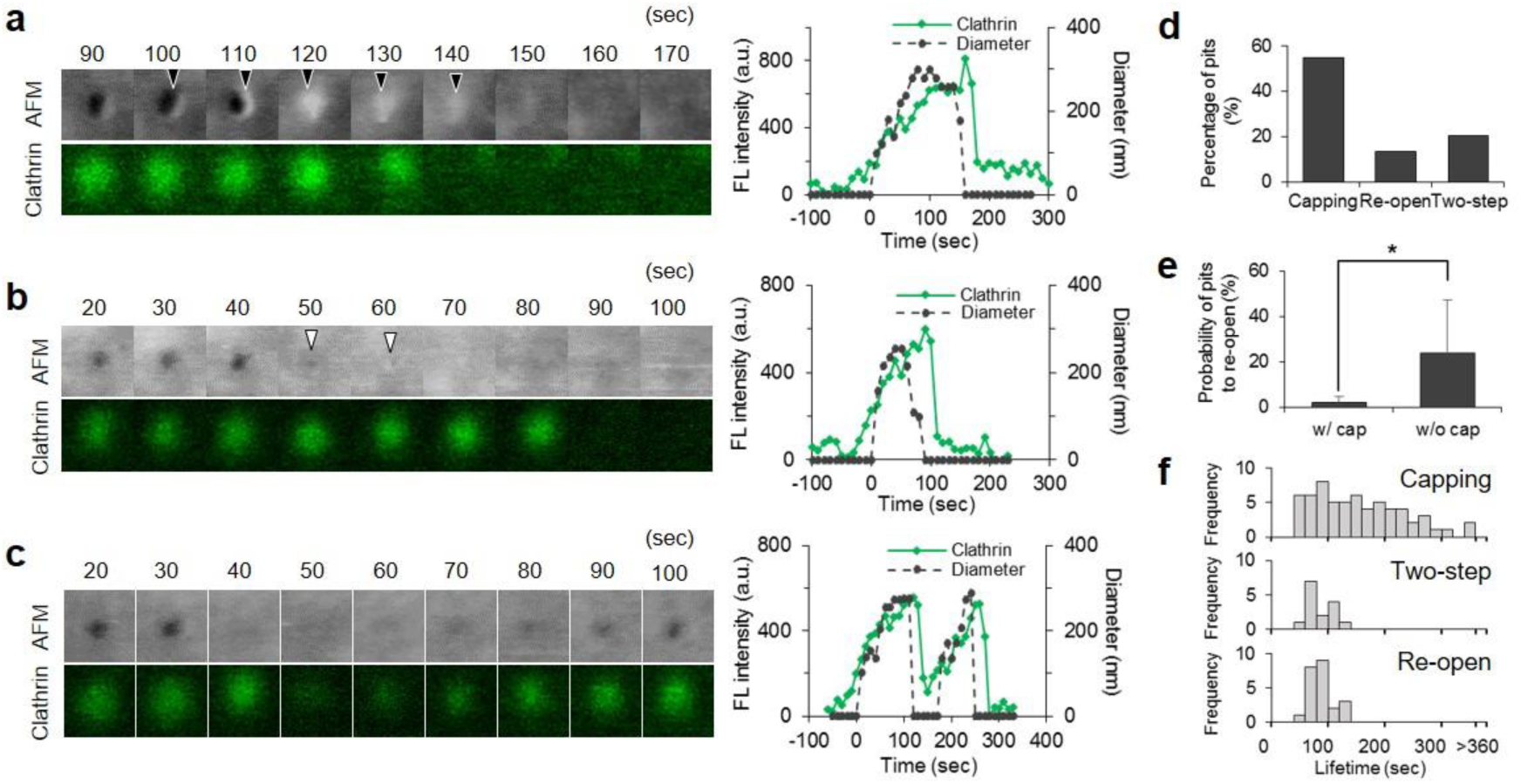
Variations in the closing motions of CCPs. Membrane morphologies of CCPs when they close were analyzed from the same data set shown in Fig. 3. (**a,b,c**) Three different motions identified in the CCPs: capping (**a**), two-step (**b**), and re-opening (**c**). AFM images and fluorescence images of EGFP-clathrin are shown in the panels at left. Fluorescence intensity (green) and the diameter of the CCP (dotted black) are also plotted in the time-course (right panels). Image size: 0.6 × 0.6 µm^2^. Membrane swelling in the capping motion, and the small aperture in the two-step motion are indicated by black and white arrowheads, respectively. (**d**) The frequency of each closing motion. In total, 113 CCPs were analyzed. The frequency of each closing motion was counted and represented as a percentage. (**e**) Relationship between capping motion and re-opening motion. Among the CCPs that showed a re-opening motion, the ratios of the CCPs with or without a capping motion are plotted. (**f**) Distribution of the CCP lifetime for three different closing motions.

There was a clear distinction between the capping and re-opening motions: pits that closed with capping did not tend to re-open (Fig. 5e), suggesting that capping plays a role in irreversible closing. In contrast, the two-step motion was not mutually exclusive to other motions, so that we sometimes observed a two-step motion that finally culminated with capping (see Supplementary Fig. S6). The comparison of the total lifetime revealed a wide distribution in capping-ended pits, whereas two-step and re-opening motions showed a narrow distribution of about 100 sec (Fig. 5f).

### Involvement of actin turnover in the closing step

Actin and actin-related proteins are also known to contribute to CCP assembly, although their exact role is not fully understood^15,30^. We previously observed and reported the dynamic turnover of the cortical actin network^31^; actin filaments are polymerized near the plasma membrane and descend into the cytoplasm. Therefore, we first examined the effect of actin inhibitors on the CME process. The analysis of the CCP lifetime in the presence of actin inhibitors revealed an inhibitory effect of the cortical actin network on the progress of CME. Cytochalasin B, an inhibitor of actin polymerization, and CK666, an inhibitor of the Arp2/3 complex, which binds to F-actin and generates a branching point, both shortened the CCP lifetime, whereas Jasplakinolide, which inhibits actin depolymerization and stabilizes the cortical actin network^31^, prolonged the lifetime (Fig. 6a, Supplementary Table 1, Supplementary Movie S5-7). In the presence of cytochalasin B, both the growing and stable phases shortened from 45 ± 34 sec to 7 ± 12 sec and 88 ± 29 sec to 59 ± 33 sec, respectively, whereas there were almost no effects on the duration of the closing step (31 ± 9 to 22 ± 7 sec) (Fig. 6b, Supplementary Table 2, Supplementary Movie S8). This indicates that the collapse of the actin network accelerates CCP assembly and maturation, whereas the stabilization of the network inhibits the process. Dissecting the two-step closing motion also revealed that cytochalasin B and CK666 shortened the duration of the large aperture, whereas Jasplakinolide prolonged it (Fig. 6a), implying that actin dynamics accelerate the assembly of CCP-related proteins.

**Figure 6.**
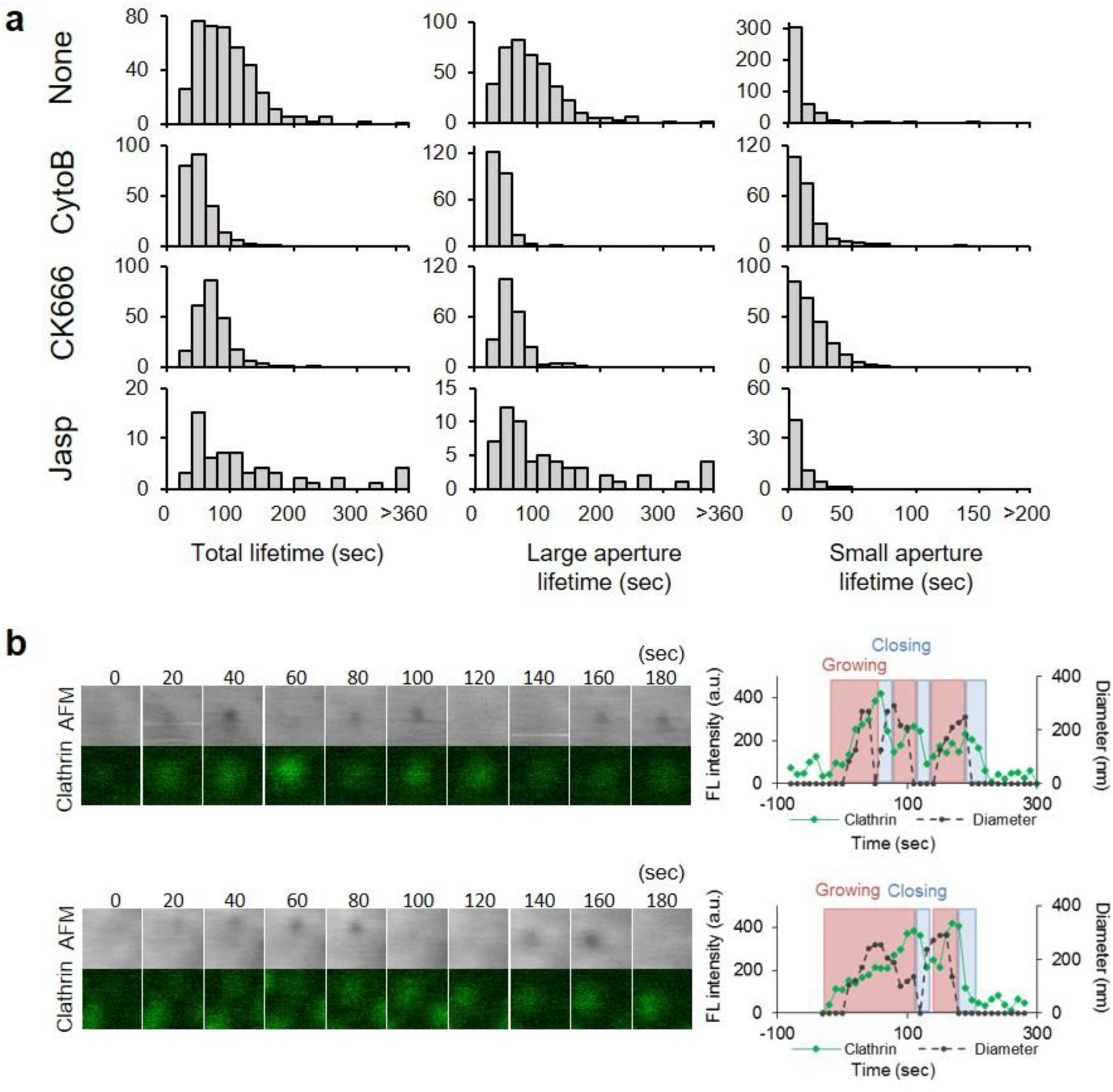
Effects of actin-related inhibitors on CCP lifetime. (**a**) COS-7 cells on the microscope stage were treated with cytochalasin B (cytoB), jasplakinolide (jasp), or CK666, and were subjected to time-lapse hybrid imaging. The total lifetime, durations of large and small apertures in the two-step motion were measured and summarized as histograms. (**b**) Time-lapse AFM and fluorescence images obtained from COS-7 cells expressing EGFP-Clathrin after treatment with cytoB (left panels). Image size: 0.6 × 0.6 µm^2^. The fluorescence intensity (green) and the diameter of the CCPs in the AFM image (dotted black) are plotted against time (right panels). The process is divided into growing (red), stable (orange), and closing (blue) phases as described in Figure 2.

In addition to the lifetime of the CCP, actin dynamics are involved in the closing motion of the CCP. The most striking effect of the inhibitors was a reduction of the capping motion and an increase in re-opening motions; cytochalasin B and CK666 drastically reduced the frequency of capping motions (from 50 to 2% by cytochalasin B, and to 5% by CK666), and increased the frequency of re-opening motions (from 20 to 70% by cytochalasin B and to 50% by CK666) (Fig. 7a, 7c). Jasplakinolide, also showed a similar effect, but to a smaller extent (Fig. 7a, 7c). These observations are in good agreement with the result that the capping and re-opening motions are inversely related (Fig. 5e), and that actin polymerization plays a role in an efficient and irreversible closing of the vesicle. In addition to the re-opening motions, two-step motions were also increased by cytochalasin B and CK666 treatments (Fig. 7a and 6a), suggesting that two-step motions and actin polymerization are tightly coupled. Blocking actin de-polymerization, but not polymerization, affected the frequency of CCP formation as previously reported^30^, implying that the cortical actin layer also has an inhibitory effect on CCP formation.

**Figure 7.**
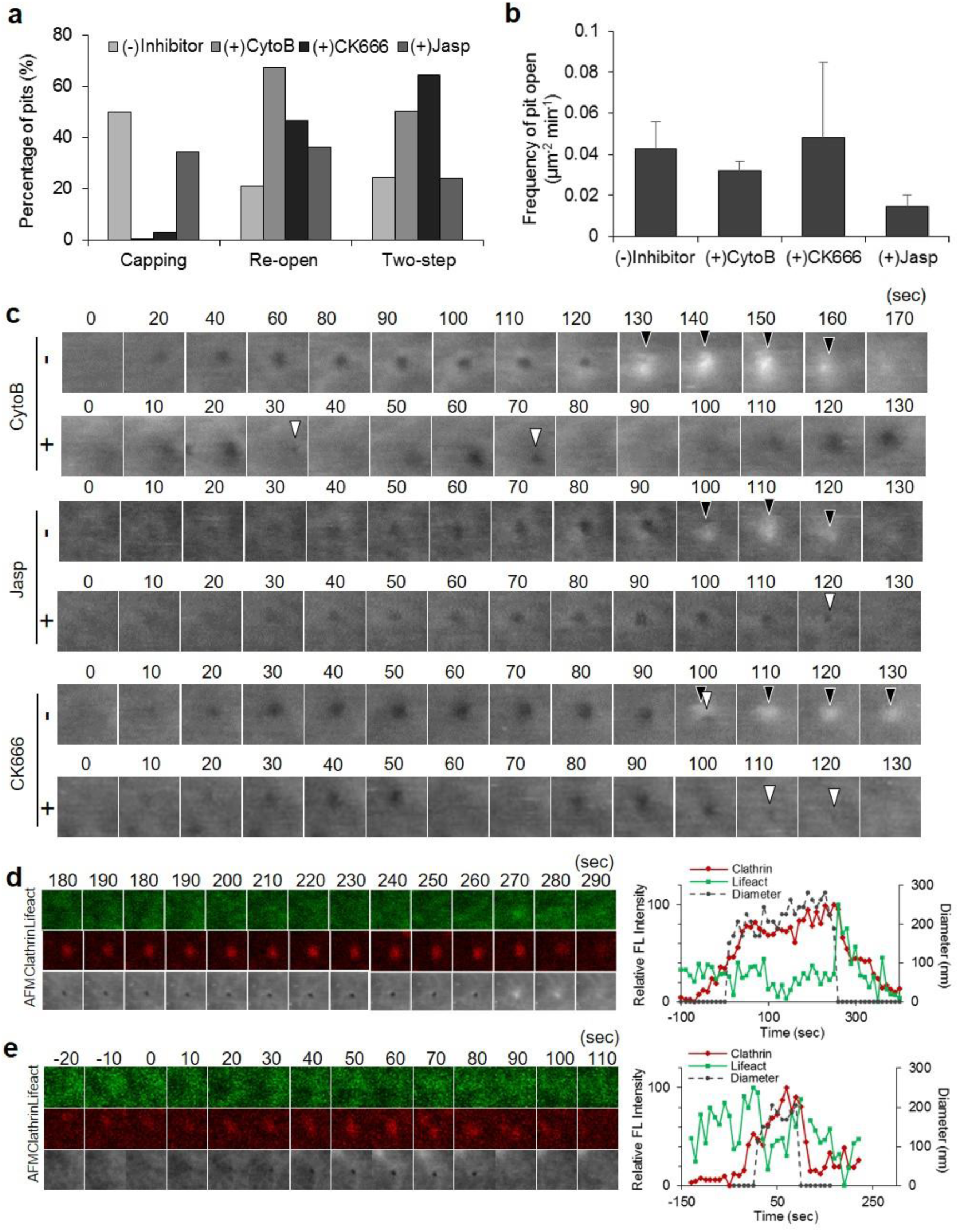
Role of actin in the closing step of CME. (**a**) The frequencies of capping, re-opening, and two-step motions were counted as described in Fig. 5d in the absence or presence of cytochalasin B (cyto B), CK666, or jasplakinolide (jasp). (**b**) The frequency of CCP formation was quantified in the absence or presence of the inhibitors and summarized. (**c**) Time-lapse AFM images obtained in a living COS-7 cell before or after treatment with CytoB, CK666 or Jasp. Image size: 0.5 × 0.5 µm^2^. Capping and the small aperture are indicated by black and white arrowheads, respectively. (**d,e**) A summary of actin assembly at a CCP from time-lapse hybrid imaging of COS-7 cells expressing EGFP-Lifeact and mCherry-clathrin. Fluorescence images for Lifeact and clathrin, as well as AFM images are shown every 10 seconds (left panels). Image size: 1.2 × 1.2 µm^2^. Two representative CCPs are shown here. The CCP closes with (**d**) and without (**e**) capping. The signal intensities of the fluorescence spots (for Lifeact (green) and clathrin (red)) and the diameter of the membrane invagination in the AFM image (black) are plotted against ti

To confirm that membrane swelling in the capping motion was induced by actin, we followed the localization of actin during CME. COS-7 cells simultaneously expressing Lifeact-GFP and mCherry-clathrin were subjected to time-lapse hybrid imaging. There were several variations in the actin signal depending on the basal level of actin around the CCP (Fig. 7d, Supplementary Movie S9). When the basal level was low, a burst of actin assembly was observed when the CCP closed. More precisely, it started to increase ~50 sec before the pit closed, and peaked slightly after (~7 sec) the pit closure (Fig. 3e, 7d,). The burst of actin signal intensity was tightly correlated with the membrane swelling in the capping motion; the actin signal peaked when the membrane swelled. This is in good agreement with the result obtained with cytochalasin B (Fig. 7a), where addition of cytochalasin B reduced the frequency of the capping motion. On the other hand, when the basal actin level around the CCP was high, the signal first decreased during the growing and stable phases, then increased again toward the end of the CME. In this case, the membrane swelling was not observed. Taken together, these results demonstrate that actin depolymerization occurs in the growth and maturation phases of CME, and active actin assembly is required for the irreversible scission of the vesicle from the plasma membrane. Furthermore, the swelling of the membrane sometimes developed into a ruffle-like protrusion, even after the pit closure (Supplementary Fig. S7), demonstrating that active polymerization of actin occurs around the CCP and generates local forces on the membrane.

### Dynamin is involved in the complete closure of CCPs

The involvement of other CCP-related proteins in morphological changes of the membrane was further investigated using RNA interference. Knockdown of dynamin2 (Fig. 8a) did not affect the frequency of pit formation (Fig. 8b), but reduced the occurrence of the capping motion from 70 to 35% (Fig. 8c, 8d. Supplementary Movie S10). The effect was similar to what was observed in the presence of cytochalasin B (Fig. 7a). However, in contrast to cytochalasin B and CK666, which increased both re-open and two-step motions, dynamin-knockdown markedly increased the two-step motion but only slightly increased the re-open motion (Fig. 8c). The detailed analysis of the two-step motion revealed a prolonged duration of the small aperture (Fig. 8e, Supplementary Table 3). These results suggest that dynamin is involved in the capping formation as well as the complete closing of the CCP.

**Figure 8.**
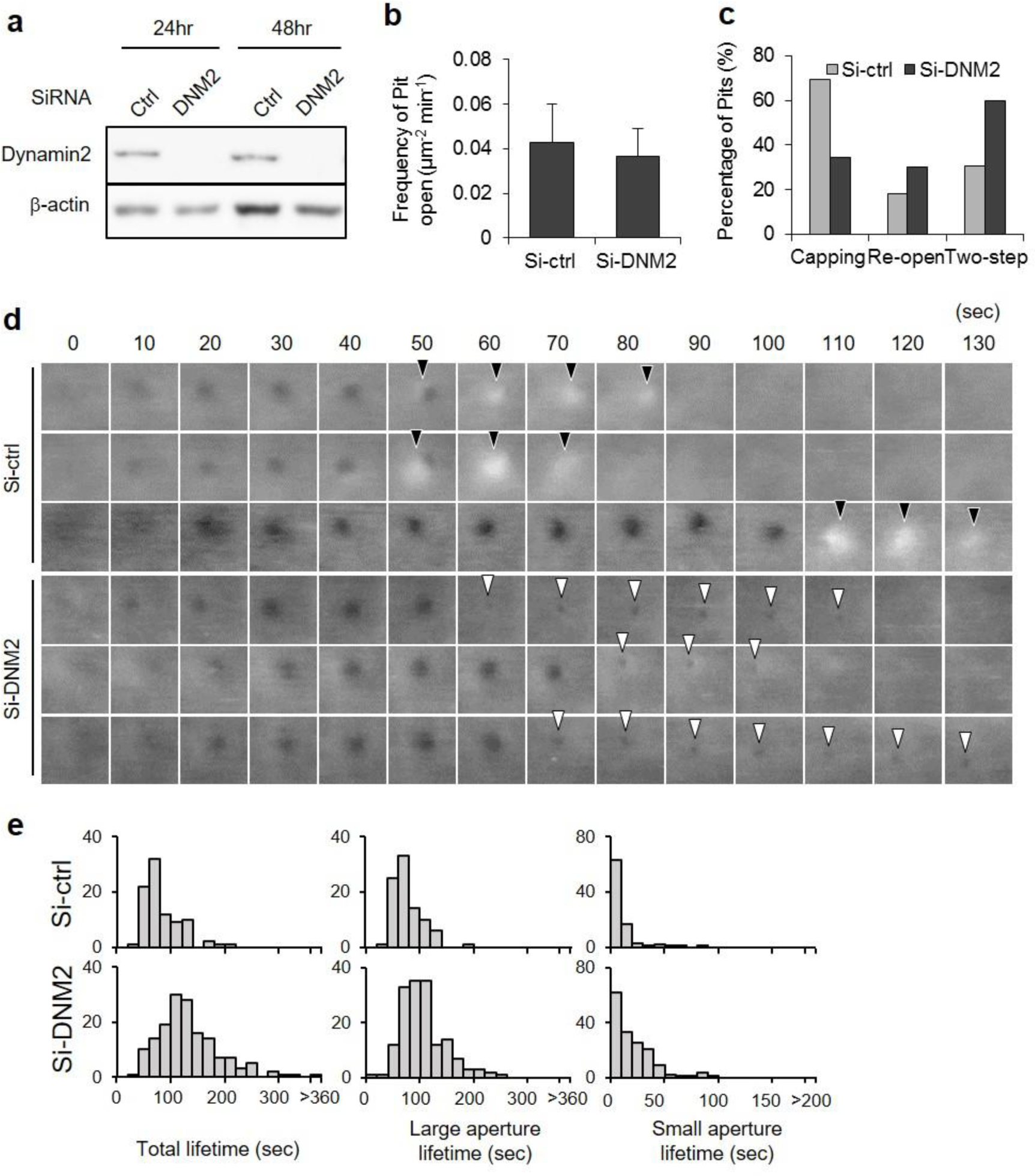
Effects of dynamin knockdown on CME. (**a**) Knockdown efficiency of dynamin2 in COS-7 cells was examined by western blotting. Total cell lysates of COS-7 cells transfected with siRNA targeting the dynamin2 gene (si-*DNM2*) or control siRNA (Luciferase, si-ctrl) were prepared 24 and 48 hours after the transfection, and were subjected to western blotting using anti-dynamin2 and β-actin antibodies. The density of the band corresponding to dynamin was quantified using β-actin as a loading control. The knockdown efficiency was estimated to be 88% after 24 hrs, and 92% after 48 hrs. (**b**) The frequency of CCP formation was counted and compared in si-ctrl and si-*DNM2* cells. (**c**) The frequency of capping, re-opening and two-step motions were analyzed in dynamin and control depleted cells. (**d**) Time-lapse AFM images obtained in a living COS-7 cell transfected with si-ctrl or si-*DNM2*. Image size: 0.5 × 0.5 µm^2^. Capping and small apertures are indicated by black and white arrowheads, respectively. (**e**) The total lifetime and the durations of large and small apertures, measured in control and dynamin-depleted cells, are summarized.

In addition to the prolonged duration of the small aperture, the duration of the large aperture was also prolonged in the dynamin-knockdown cells, resulting in the lengthening of the total lifetime from 61 ± 26 sec (n = 90) to 80 ± 31 sec (n =111) (Fig. 8e, Supplementary Table 3). This is in good agreement with our observation that dynamin started to appear at the CCP during the growing phase (Fig. 3c, 3d). These results suggested that dynamin is involved not only in the closing step, but also in the assembly and maturation phases of the CCP, as was suggested in a previous study^24^. It should be noted that dynamin knockdown did not completely block the progress of the CME, implying that dynamin is not absolutely necessary for CME.

## Discussion

Imaging the shape of the plasma membrane has been technically challenging. Although AFM was first applied to a living cell in 1992, its low scanning rate did not allow visualization of the endocytic process. Many other microscopic techniques such as high-frequency microrheology^32^ and high-speed ion conductance microscopy^33^ have also been utilized for cell surface imaging and characterization. In this study, we visualized morphological changes of the plasma membrane during CME in a living cell using HS-AFM coupled with confocal laser-scanning microscopy, and revealed unique membrane structures at the end stage of CME as well as the role of CME-related proteins in such morphological dynamics. To our knowledge, this is the first direct observation of the CME process without labeling, fixation, or staining. The innovation of the hybrid microscope system is that the localization of a specific protein of interest can be mapped on a series of AFM images.

Recent advances in fluorescence microscopy enabled highly-precise spatiotemporal analyses of proteins involved in the CME in yeast^34,35^ and animal cells^5^. They revealed the timings of protein assembly at the CCP and disassembly after the pit closure. On the other hand, the morphological changes of the plasma membrane during the CME were mainly drawn based on EM snapshots of fixed and stained cells. The localization of a specific protein on the CCP has also been revealed by immune-electron microscopy and by fluorescence-electron microscopy. For example, clathrin exists at the place where the membrane is slightly bent^12,13^. However, the time-resolution of the snapshot analysis is limited, and is not suitable for following membrane dynamics. In time-lapse fluorescence imaging, a complete pit closure was detected by a pH-sensitive fluorescent dye combined with fast exchange of the external medium between high and low pH solutions^5,36,37^, but other morphological changes of the membrane, such as invagination and protrusion, could not be detected. Our time-lapse hybrid imaging of HS-AFM and CLSM rendered both the membrane morphology and protein localization with a time resolution of several seconds, which was particularly suitable for tracking the morphological changes of the CCP together with the assembly of specific proteins.

Our image analysis of hundreds of CMEs revealed several unique membrane dynamics at the end of the process (Fig. 5): capping, a two-step motion, and re-opening. Capping occurred in most of the CME events (~70%) and was mediated by actin (Fig. 7) and dynamin (Fig. 8). A region of the adjacent membrane swelled and covered over the CCP (Fig. 5, 7). The peak actin signal (GFP-Lifeact) corresponded with the timing of membrane swelling (Fig. 7d), and the inhibition of actin polymerization by inhibitors (cytochalasin B and CK666) perturbed the capping motion (Fig. 7a), indicating that the membrane swelling is caused by rapid and local actin polymerization. Notably, the inhibitor not only decreased the frequency of capping events, but also increased the frequency of re-opening events (Fig. 7a). This clearly indicates that without cortical actin dynamics, the membrane vesicle could reversibly fuse back to the plasma membrane. It is still unclear whether the vesicle is completely detached from the membrane, or is still docked via a protein tether. However, our observation that the clathrin signal diminished after the first closure and then increased upon re-opening (Fig. 5c, Fig. 6b) suggests that the CCP machinery is somehow disassembled during the closed state.

The role of actin dynamics in CME has been debated in previous studies^38,39^. The cortical actin layer is known to be actively involved not only in endocytosis but also exocytosis. In a secretion process of neuronal cells, the cortical actin network is involved in (i) tethering secretory vesicles^40–45^, (ii) providing a platform for directed movement toward the plasma membrane^46^ and (iii) facilitating the generation of new release sites^47–50^. In endocytosis, actin is known to localize to the CCP in both yeast and mammalian cells^34,35^. Although several models have been proposed for the function of actin in CME, many details remain obscure. Our observation that actin inhibitors drastically decreased the capping motion and increased the occurrence of re-opening motions (Fig. 7a) suggests that actin dynamics are required for irreversible detachment of the CCV from the plasma membrane. In one possible scenario, actin could polymerize around the vesicle and provide a driving force to push the vesicle into the cytoplasm by interacting with myosin in the cell cortex. There are a number of reports of non-muscle myosins in the cell cortex^51–53^, which are involved in various molecular events at the cell surface. Another possibility is that actin assembles near the plasma membrane but does not attach to the vesicle. Newly assembled actin filaments between the plasma membrane and the vesicle may spatially hinder the reversible fusion of the vesicle to the plasma membrane, which consequently pushes the vesicle into the cytoplasm, ultimately leading to endosome fusion, as was suggested by previous studies^39,54^. It is intriguing that most of the membrane swelling occurs at one side of the CCP and moves across the pit towards the opposite side (Fig. 5, 7). We could not find any preference in the direction of the capping (not shown, or suppl.). It might be the case that a sudden burst of actin polymerization at a certain point on the CCP induces membrane swelling.

In addition to the closing motion, we found a role of actin dynamics in the growing phase of the CCP. Our observation that destruction of the actin network by cytochalasin B or CK666 accelerated the CME process (shortened the lifetime of open apertures), and stabilization of the network by Jasplakinolide prolonged the lifetime (Fig. 6) suggests that the cortical actin layer has an inhibitory effect on the progress of the CME. This could be explained by the following two mechanisms. The first possibility is that the assembly of CCP protein components at the plasma membrane is inhibited by the cortical actin layer. Since the cortical layer is supposed to be similar to a hydrogel state, the diffusion of cytoplasmic proteins through the cortex is also supposed to be slow. The second possibility is that the cortical actin layer spatially inhibits the growth of the CCP. As the size of the CCP grows, neighboring actin filaments must be excluded. Although it is not clear whether the exclusion is mediated by physical force or an enzymatic process^55,56^, the progress of the CME is tightly coupled with the dynamics of the cortical actin network.

In addition to actin, we found that dynamin plays a role in the closing motion (Fig. 8). Knocking down of dynamin reduced the capping and re-opening motion, and increased the two-step motion (Fig. 8). The loss of dynamin apparently prolonged the duration of a smaller aperture size (Fig. 8), suggesting that dynamin plays a role not in the initial narrowing of the aperture, but in the complete scission of the vesicle. This is compatible with the known function of dynamin: it binds to the neck of the CCP and catalyzes membrane scission^57,58^. On the other hand, a recent *in vitro* study reported that actin and BAR domain proteins, but not dynamin, are essential for membrane scission^59^. Therefore, it might be the case that the initial narrowing of the pit aperture is mediated by actin and BAR domain proteins, and the final scission step might be accelerated by a catalytic function of dynamin. In such a case, the inhibition of individual proteins would not completely abolish the complete process but would only slow down the progress.

We found that knocking down dynamin also prolonged the assembly and/or maturation phases of CME (Fig. 8). Indeed, dynamin appeared on the CCP just prior to the initiation of membrane invagination, and kept accumulating as the pit grew (Fig. 3)^60^. The role of dynamin in the assembly and maturation phases has been debated in previous studies^6,24^. Since the catalytic activity of dynamin is regulated by nucleotide-based mechanisms^58^, the assembly of dynamin at the CCP may not in itself induce any membrane deformations. Considering the fact that the dynamin knockdown slowed down the progress of CME, it might be the case that dynamin is necessary for recruiting other protein machineries to the CCP. Further analyses are required for elucidating the role of dynamin in the whole process of CME.

## Methods

### Materials

COS-7 cells were purchased from DS Biopharma Medical (EC87021302-F0). Cytochalasin B was purchased from Sigma-Aldrich (St. Louis, USA), and jasplakinolide was purchased from Abcam (Cambridge, UK). The reagents were added to the culture medium at final concentrations of 2 μM for cytochalasin B and 1 μM for jasplakinolide. HEPES-NaOH (pH 7.0–7.6), which was used to maintain a constant pH of the medium during AFM observation, was purchased from Sigma-Aldrich. The mammalian expression vectors encoding Dyn2-pmCherryN1 and Epsin2-pmCherryC1 were gifts from Christien Merrifield (Addgene # 27689 and # 27673, respectively), and the vector for EGFP-Lifeact expression was a kind gift from Dr. Mineko Kengaku (Kyoto University, Kyoto, Japan). Silencer select siRNA targeting *DNM2* (#s4212) and siRNA targeting Luciferase (#12935-146) were purchased from Ambion and ThermoFisher Scientific (Waltham, USA), respectively. Transfection reagents Lipofectamin2000 and Effectene were purchased from Thermo Fisher Scientific (Waltham, USA) and Qiagen (Hilden, Germany), respectively. Anti-dynamin2 antibody was from Cell Signaling Technology (Danvers, USA), and PVDF membrane was from Bio-Rad Laboratories (Hercules, USA).

### Cell culture, transfection and fixation

One or two days before AFM imaging, COS-7 monkey kidney derived fibroblast-like cells were seeded on a poly-L-lysine-coated glass slide and grown at 37°C with 5% CO2 in Dulbecco’s Modified Eagle’s Medium (DMEM) supplemented with 10% fetal bovine serum (FBS). AFM imaging was performed in DMEM supplemented with 10% FBS and 10 mM HEPES-NaOH (pH 7.0–7.6). cDNAs encoding human clathrin light chain A (CLTA, NM_007096) and caveolin 1 (CAV1, NM_00172895) were amplified by RT-PCR and cloned into the vector pEGFP-C3 (Clontech) to create fused proteins with enhanced green fluorescent protein (EGFP). The plasmids were introduced into cells using Effectene Transfection Reagent according to the manufacturer’s protocol. At 24–48 h after transfection, the cells were used for AFM imaging. Expression of the fusion protein was confirmed by fluorescence signals from the cells. For experiments with fixed cells, the cells were fixed with 5% paraformaldehyde in PBS for 15 min at room temperature and washed with PBS.

### RNA interference

Cells were transfected with siRNAs using Lipofectamine 2000 (Invitrogen) following the manufacturer’s instructions, and were harvested after 24-48 hours for analysis by SDS-PAGE and immunoblotting, using PVDF membranes. Each membrane was cut into two halves at the protein size of 60 kDa; the upper half was blotted with anti-dynamin2 antibody, and the bottom half was detected with anti-β-actin antibody.

### AFM imaging and data analysis

BIXAM™ (Olympus Corp., Tokyo, Japan), which is a tip-scan type HS-AFM unit combined with an inverted fluorescent/ optical microscope (IX83, Olympus) equipped with a phase contrast system and a confocal unit (FV1200, Olympus), was used for this study. The tip-scan HS-AFM imaging system was developed based on a previous study^21^. In brief, the modulation method was set to phase modulation mode to detect tip-sample interactions. An electron-beam deposited sharp cantilever tip with a spring constant of 0.1 N m^−1^ (USC-F0.8-k0.1, a customized cantilever from Nanoworld [Neuchâtel, Switzerland]) was used. The AFM tip was aligned with confocal views as described in the Results section. The images from the confocal microscope and AFM were simultaneously acquired at a scanning rate of 0.1 frames per second. The captured sequential images were overlaid by using AviUTL (http://spring-fragrance.mints.ne.jp/aviutl/) based on the tip position. The fluorescence intensity was quantified by Image J software, (http://rsbweb.nih.gov/ij/). The lifetime of the pit was analyzed with Metamorph imaging software (Molecular Devices). The diameter/height of the pit or membrane swelling region was obtained using AFM Scanning System Software Version 1.6.0.12 (Olympus).

### Statistical analysis

Data presented as graphs are from three independent experiments. The number of total CCPs analyzed for each analysis is specified in figure legends. Statistical analysis was performed by two-way analysis of variance followed by Student’s t-test.

## Acknowledgements

This study was supported by a Grant-in-Aid for Challenging Exploratory Research (No. 16K14722 to S.H.Y.) from the Japan Society for the Promotion of Science (JSPS), and the Advanced Research & Development Programs for Medical Innovation from the Japan Agency for Medical Research and Development (No. 16gm5810018h0001 to S.H.Y.). A.Y. is a recipient of a JSPS research fellowship. Support from Building of Consortia for the Development of Human Resources in Science and Technology to Y.S. is also acknowledged. We would also like to thank Mr. Shuichi Ito and Mr. Akira Yagi for their technical assistance.

## Author contributions

A.Y., N.S., and S.H.Y. conceived and designed the experiments on living cells. N.S., Y.U., and Y.Imaoka developed the combined CLSM-high-speed-AFM instrument. A.Y., N.S., Y.U., and Y. Itagaki performed experiments. A.Y., N.S., Y.Itagaki and S.H.Y. analyzed the data. A.Y., S.N., Y.S., Y. Itagaki and S.H.Y. co-wrote the manuscript. All authors discussed the results and commented on the manuscript.

## References

1. Boucrot, E. et al. Endophilin-A2 functions in membrane scission in clathrin-independent endocytosis’. (2014). doi:10.1038/nature14064

2. Doherty, G. J. & McMahon, H. T. Mechanisms of endocytosis. Annu. Rev. Biochem. 78, 857–902 (2009).

3. Hansen, C. G. & Nichols, B. J. Molecular mechanisms of clathrin-independent endocytosis. J. Cell Sci. 122, 1713–21 (2009).

4. McMahon, H. T. & Boucrot, E. Molecular mechanism and physiological functions of clathrin-mediated endocytosis. Nat. Rev. Mol. Cell Biol. 12, 517–533 (2011).

5. Taylor, M. J., Perrais, D. & Merrifield, C. J. A high precision survey of the molecular dynamics of mammalian clathrin-mediated endocytosis. PLoS Biol. 9, (2011).

6. Taylor, M. J., Lampe, M. & Merrifield, C. J. A feedback loop between dynamin and actin recruitment during clathrin-mediated endocytosis. PLoS Biol. 10, (2012).

7. Merrifield, C. J., Feldman, M. E., Wan, L. & Almers, W. Imaging actin and dynamin recruitment during invagination of single clathrin-coated pits. Nat Cell Biol 4, 691–698 (2002).

8. Jarsch, I. K., Daste, F. & Gallop, J. L. Membrane curvature in cell biology: An integration of molecular mechanisms. J. Cell Biol. 214, 375–387 (2016).

9. Dawson, J. C., Legg, J. A. & Machesky, L. M. Bar domain proteins: a role in tubulation, scission and actin assembly in clathrin-mediated endocytosis. Trends Cell Biol. 16, 493–498 (2006).

10. Suetsugu, S. Higher-order assemblies of BAR domain proteins for shaping membranes. Microscopy 65, 201–210 (2016).

11. Daumke, O., Roux, A. & Haucke, V. BAR domain scaffolds in dynamin-mediated membrane fission. Cell 156, 882–892 (2014).

12. Avinoam, O., Schorb, M., Beese, C. J., Briggs, J. A. G. & Kaksonen, M. Endocytic sites mature by continuous bending and remodeling of the clathrin coat. Science 348, 1369–72 (2015).

13. Sochacki, K. A., Dickey, A. M., Strub, M.-P. & Taraska, J. W. Endocytic proteins are partitioned at the edge of the clathrin lattice in mammalian cells. Nat. Cell Biol. 19, 352–361 (2017).

14. Park, R. J. et al. Dynamin triple knockout cells reveal off target effects of commonly used dynamin inhibitors. J. Cell Sci. 126, 5305–12 (2013).

15. Ferguson, S. M. et al. Coordinated actions of actin and BAR proteins upstream of dynamin at endocytic clathrin-coated pits. Dev. Cell 17, 811–22 (2009).

16. Takei, K., McPherson, P. S., Schmid, S. L. & De Camilli, P. Tubular membrane invaginations coated by dynamin rings are induced by GTP-gamma S in nerve terminals. Nature 374, 186–90 (1995).

17. Kodera, N., Yamamoto, D., Ishikawa, R. & Ando, T. Video imaging of walking myosin V by high-speed atomic force microscopy. Nature 468, 72–6 (2010).

18. Yokokawa, M. et al. Fast-scanning atomic force microscopy reveals the ATP/ADP-dependent conformational changes of GroEL. EMBO J. 25, 4567–4576 (2006).

19. Igarashi, K. et al. Traffic Jams Reduce Hydrolytic Efficiency of Cellulase on Cellulose Surface. Science (80-.). 333, 1279–1282 (2011).

20. Colom, A., Redondo-Morata, L., Chiaruttini, N., Roux, A. & Scheuring, S. Dynamic remodeling of the dynamin helix during membrane constriction. Proc. Natl. Acad. Sci. U. S. A. 114, 5449–5454 (2017).

21. Suzuki, Y. et al. High-speed atomic force microscopy combined with inverted optical microscopy for studying cellular events. Sci. Rep. 3, (2013).

22. Yoshida, A. et al. Probing in vivo dynamics of mitochondria and cortical actin networks using high-speed atomic force/fluorescence microscopy. Genes to Cells 20, 85–94 (2015).

23. Avinoam, O., Schorb, M., Beese, C. J., Briggs, J. A. G. & Kaksonen, M. Endocytic sites mature by continuous bending and remodeling of the clathrin coat. Science 348, 1369–72 (2015).

24. Loerke, D. et al. Cargo and Dynamin Regulate Clathrin-Coated Pit Maturation. PLoS Biol. 7, e57 (2009).

25. Lee, D.-W., Wu, X., Eisenberg, E. & Greene, L. E. Recruitment dynamics of GAK and auxilin to clathrin-coated pits during endocytosis. J. Cell Sci. 119, 3502–3512 (2006).

26. Greener, T., Zhao, X., Nojima, H., Eisenberg, E. & Greene, L. E. Role of cyclin G-associated kinase in uncoating clathrin-coated vesicles from non-neuronal cells. J. Biol. Chem. 275, 1365–1370 (2000).

27. Horvath, C. a J., Vanden Broeck, D., Boulet, G. a V, Bogers, J. & De Wolf, M. J. S. Epsin: Inducing membrane curvature. Int. J. Biochem. Cell Biol. 39, 1765–1770 (2007).

28. Aguet, F., Antonescu, C. N., Mettlen, M., Schmid, S. L. & Danuser, G. Advances in analysis of low signal-to-noise images link dynamin and AP2 to the functions of an endocytic checkpoint. Dev. Cell 26, 279–291 (2013).

29. Stan, R. V. Structure of caveolae. Biochim. Biophys. Acta – Mol. Cell Res. 1746, 334–348 (2005).

30. Grassart, A. et al. Actin and dynamin2 dynamics and interplay during clathrin-mediated endocytosis. J. Cell Biol. 205, 721–735 (2014).

31. Zhang, Y. et al. In vivo dynamics of the cortical actin network revealed by fast-scanning atomic force microscopy. Reprod. Syst. Sex. Disord. 1–11 (2017). doi:10.1093/jmicro/dfx015

32. Rigato, A., Miyagi, A., Scheuring, S. & Rico, F. High-frequency microrheology reveals cytoskeleton dynamics in living cells. Nat. Phys. 13, (2017).

33. Shevchuk, A. I. et al. An alternative mechanism of clathrin-coated pit closure revealed by ion conductance microscopy. J. Cell Biol. 197, 499–508 (2012).

34. Toshima, J. Y. et al. Spatial dynamics of receptor-mediated endocytic trafficking in budding yeast revealed by using fluorescent-factor derivatives. Proc. Natl. Acad. Sci. 103, 5793–5798 (2006).

35. Kaksonen, M., Toret, C. P. & Drubin, D. G. A modular design for the clathrin- and actin-mediated endocytosis machinery. Cell 123, 305–320 (2005).

36. Kirchhausen, T. Imaging endocytic clathrin structures in living cells. Trends Cell Biol. 19, 596–605 (2009).

37. Merrifield, C. J., Perrais, D. & Zenisek, D. Coupling between clathrin-coated-pit invagination, cortactin recruitment, and membrane scission observed in live cells_Supplenetal information. Cell 121, 593–606 (2005).

38. Fujimoto, L. M., Roth, R., Heuser, J. E. & Schmid, S. L. Actin assembly plays a variable, but not obligatory role in receptor-mediated endocytosis in mammalian cells. Traffic 1, 161–171 (2000).

39. Boulant, S., Kural, C., Zeeh, J.-C., Ubelmann, F. & Kirchhausen, T. Actin dynamics counteract membrane tension during clathrin-mediated endocytosis. Nat. Cell Biol. 13, 1124–1131 (2011).

40. Aschenbrenner, L., Lee, T. & Hasson, T. Myo6 facilitates the translocation of endocytic vesicles from cell peripheries. Mol. Biol. Cell 14, 2728–2743 (2003).

41. Chibalina, M. V, Seaman, M. N. J., Miller, C. C., Kendrick-Jones, J. & Buss, F. Myosin VI and its interacting protein LMTK2 regulate tubule formation and transport to the endocytic recycling compartment. J. Cell Sci. 120, 4278–4288 (2007).

42. Desnos, C., Huet, S. & Darchen, F. ‘Should I stay or should I go?’: myosin V function in organelle trafficking. Biol. cell 99, 411–23 (2007).

43. Tomatis, V. M. et al. Myosin vi small insert isoform maintains exocytosis by tethering secretory granules to the cortical actin. J. Cell Biol. 200, 301–320 (2013).

44. Huet, S. et al. Myrip Couples the Capture of Secretory Granules by the Actin-Rich Cell Cortex and Their Attachment to the Plasma Membrane. J. Neurosci. 32, 2564–2577 (2012).

45. Arden D. S., Puri, C., Au, J. S.-Y., Kendrick-Jones, J. & Buss, F. Myosin VI Is Required for Targeted Membrane Transport during Cytokinesis. Mol. Biol. Cell 18, 4750–4761 (2007).

46. Papadopulos, A. et al. Activity-driven relaxation of the cortical actomyosin II network synchronizes Munc18-1-dependent neurosecretory vesicle docking. Nat. Commun. 6, 6297 (2015).

47. Papadopulos, A., Tomatis, V. M., Kasula, R. & Meunier, F. a. The Cortical Acto-Myosin Network: From Diffusion Barrier to Functional Gateway in the Transport of Neurosecretory Vesicles to the Plasma Membrane. Front. Endocrinol. (Lausanne). 4, 153 (2013).

48. de Paiva, A., Meunier, F. A., Molgó, J., Aoki, K. R. & Dolly, J. O. Functional repair of motor endplates after botulinum neurotoxin type A poisoning: biphasic switch of synaptic activity between nerve sprouts and their parent terminals. Proc. Natl. Acad. Sci. U. S. A. 96, 3200–5 (1999).

49. Zakharenko, S., Chang, S., O’Donoghue, M. & Popov, S. V. Neurotransmitter secretion along growing nerve processes: Comparison with synaptic vesicle exocytosis. J. Cell Biol. 144, 507–518 (1999).

50. Papadopulos, A. Membrane shaping by actin and myosin during regulated exocytosis. Mol. Cell. Neurosci. (2017). doi:10.1016/j.mcn.2017.05.006

51. Chandrasekar, I. et al. Nonmuscle myosin II is a critical regulator of clathrin-mediated endocytosis. Traffic 15, 418–432 (2014).

52. Buss, F., Luzio, J. P. & Kendrick-Jones, J. Myosin VI, a new force in clathrin mediated endocytosis. FEBS Lett. 508, 295–299 (2001).

53. Cheng, J., Grassart, A. & Drubin, D. G. Myosin 1E coordinates actin assembly and cargo trafficking during clathrin-mediated endocytosis. Mol. Biol. Cell 23, 2891–904 (2012).

54. Collins, A., Warrington, A., Taylor, K. a. & Svitkina, T. Structural organization of the actin cytoskeleton at sites of clathrin-mediated endocytosis. Curr. Biol. 21, 1167–1175 (2011).

55. Engqvist-Goldstein, A. E. Y. et al. RNAi-mediated Hip1R silencing results in stable association between the endocytic machinery and the actin assembly machinery. Mol. Biol. Cell 15, 1666–79 (2004).

56. Clainche, C. Le, Pauly, B. S., Zhang, C. X., Cunningham, K. & Drubin, D. G. A Hip1R – cortactin complex negatively regulates actin assembly associated with endocytosis. 26, 1199–1210 (2007).

57. Morlot, S. & Roux, A. Mechanics of dynamin-mediated membrane fission. Annu. Rev. Biophys. 42, 629–49 (2013).

58. Antonny, B. et al. Membrane fission by dynamin: what we know and what we need to know. EMBO J. 35, 2270–2284 (2016).

59. Simunovic, M. et al. Friction Mediates Scission of Tubular Membranes Scaffolded by BAR Proteins. Cell 172–184 (2016). doi:10.1016/j.cell.2017.05.047

60. Damke, H., Binns, D. D., Ueda, H., Schmid, S. L. & Baba, T. Vesicle Formation at Morphologically Distinct Stages. Mol. Biol. Cell 12, 2578–2589 (2001).

61. Shevchuk, A. I. et al. An alternative mechanism of clathrin-coated pit closure revealed by ion conductance microscopy. J. Cell Biol. 197, 499–508 (2012).

